# Selected for the surface: adaptive laboratory evolution for accelerated cathodic biofilm development in *Cupriavidus necator*

**DOI:** 10.64898/2026.11.09.765286

**Authors:** Sri Sannihita Chavali, Yitian Zhou, Jan Hendrick Nissen, Chistian Jonas Lapp, Janek R. Weiler, Maike Orth, Stefan Heinrich, Alexandra von Kameke, Miriam Edel-Teichmann, Johannes Gescher

## Abstract

Cupriavidus necator is a metabolically versatile facultative chemolithoautotroph and a promising chassis for sustainable bioproduction from CO_2_. Efficient coupling of its metabolism to renewable electricity through microbial electrosynthesis (MES) requires the organism to form stable, electroactive biofilms on cathodes. This step is hindered under oxic conditions (oxic MES, oMES) by electrochemically generated reactive oxygen species (ROS), mass-transfer limitations and poor biofilm stability. Systematic strategies to improve *C. necator* biofilm formation on cathodes have been lacking. Here we used adaptive laboratory evolution (ALE) to address this gap. The progenitor strain colonised graphite cathodes only after an initial heterotrophic phase and formed spatially uneven biofilms whose distribution mirrored the local flow field, as resolved by micro-particle tracking velocimetry. Populations were then selected over five passages under controlled, oxygen-limited cathodic conditions. Resequencing revealed that the recovered population was genetically heterogeneous, and its dominant isolate, strain 890, was characterised in a microfluidic oMES platform. Strain 890 reached the target electrode coverage in about one day instead of six and maintained a robust biofilm at the progenitor’s steady-state biovolume, whereas the multi-strain pool peaked higher but proved unstable. Relative to the progenitor, the accelerated biofilm phenotype of strain 890 is driven by a single novel mutation in a bifunctional diguanylate cyclase/phosphodiesterase locus, which likely encodes a central regulator of the planktonic-to-biofilm transition. The steady-state biovolume itself remained bounded most probably due to a mass-transfer ceiling set by the counter-directional supply of H₂ and O₂. This work shows that ALE renders the initial-attachment barrier of cathodic biofilm formation genetically tractable, providing strain 890 as a robust chassis and a well-defined starting point for oMES-based CO₂ valorisation.

## 1. Introduction

*Cupriavidus necator* is a facultative chemolithoautotroph that grows heterotrophically on organic substrates or autotrophically using CO₂ as the sole carbon source and H₂ as energy donor via the Calvin–Benson–Bassham cycle, making it an attractive chassis organism for sustainable biomanufacturing from gaseous feedstocks (Panich et al., 2021). Autotrophic and heterotrophic cultivation strategies have been widely explored to convert CO_2_, with H_2_ as the electron donor, into biomass and value-added products including acetoin, 2,3-butanediol, terpenes and biofuels (Windhorst and Gescher, 2019; Panich et al., 2021; Weiler et al., 2024), as well as polyhydroxyalkanoates (Nangle *et al*., 2020).

Coupling this metabolism to renewable electricity through microbial electrosynthesis (MES) offers a promising route to electrify CO₂ fixation, with a cathode serving directly or indirectly as electron and energy source for electroautotrophic growth (Sydow et al., 2017; Reiner et al., 2018, 2020; Jourdin et al., 2020; Jourdin and Burdyny, 2021; J. Weiler et al., 2024; Rominger et al., 2025).

Still most literature examples of MES are still based on the activity of anaerobic methanogens and acetogens. The outstanding advantage of these organisms is their electron efficiency as the biotechnological end product is at the same time the catabolic product of these organisms, generated via the very efficient Wood-Ljungdahl CO_2_ fixation pathway (Reiner *et al*., 2020). Still energetic limitations, limit genetic development of these organisms for a broader spectrum of end products and their obligate anaerobic metabolism necessitates a strict separation between anode and cathode as the anode reaction leads to oxygen evolution. Regardless of aerobic or anaerobic nature of the bacteria, the biocatalyst can either be grown as a biofilm on the cathode or planktonic in the media and supplied with electrochemically generated H_2_. Both processes can reach high electron recoveries, and values approaching unity have been reported for planktonic cultures producing acetate or methane (Kracke *et al*., 2019). These recoveries rest, however, on the advantage outlined above, namely that in acetogens and methanogens the product of interest is at the same time the catabolic end product of the Wood-Ljungdahl pathway, so that almost every electron delivered to the cell is recovered in it. They therefore do not transfer to biocatalysts whose product is not the terminal electron sink of their metabolism. For such organisms the configuration is instead governed by the volumetric conversion rate, by the energy demand of the conversion and by the retention of the biocatalyst. A high cell density directly at the electrode surface raises the catalyst concentration per unit cathode volume well above that attainable in suspension, and the highest volumetric CO-conversion rates so far reported for MES have been achieved with dense biofilms on structured cathodes (Cabau-Peinado *et al*., 2024). Beyond simply accumulating cells, biofilms continuously consume H directly from the cathode (Jourdin *et al*., 2016; Rominger *et al*., 2025). This lowers the energy demand compared to gas fermentation, where an external electrolyser is required. The advantages extend beyond energy savings as direct H consumption overcomes its low aqueous solubility, which otherwise limits space-time yields and necessitates energy-intensive mixing. Furthermore, biofilms prevent cell washout, enabling continuous operation regardless of hydraulic retention time. This is the rationale behind the widespread use of cathodic biofilms as biocatalysts (Flexer and Jourdin, 2020; Philipp *et al*., 2024; Weiler *et al*., 2024). This benefit, however, has an inherent ceiling. As soon as H bubbles grow large enough, they shear the biofilm off the electrode and cap the attainable performance (Rominger *et al*., 2025).

Oxic electroautotrophic biofilms have been described for a small range of cathodic systems, in which microbial communities or isolated Knallgas (hydrogen-oxidizing) bacteria form dense films that couple cathodic electrons to CO₂ fixation and product formation (Reiner *et al*., 2020; Pous *et al*., 2023; Chavali *et al*., 2026). A well-characterised example is the thermoacidophilic Knallgas bacterium *Kyrpidia spormannii* EA-1, whose adaptation to cathodic growth yielded higher biofilm accumulation rates and faster and more homogeneous electrode coverage (Jung *et al*., 2020).

A defining constraint of MES operated under oxic conditions (oxic MES, oMES) is the electrochemical generation of reactive oxygen species (ROS) at the cathode. At the strongly reducing potentials typically applied (−0.3 to −0.8 V vs. standard hydrogen electrode (SHE)), dissolved O₂ can be reduced to superoxide (O₂·⁻) and hydrogen peroxide (H₂O₂) (Wang et al., 2021; Chen et al., 2023). These ROS impose oxidative stress on nascent biofilms driving lipid peroxidation, protein damage and DNA lesions that inhibits initial attachment, suppresses extracellular polymeric substance (EPS) production and triggers cell death, thereby preventing uniform colonization and biofilm maturation.

For organisms thriving via oMES, this oxidative burden will not act alone. Even where the organisms attach readily to gas-transferring electrodes, sustained productivity is limited by the combination of persistent ROS exposure and internal substrate diffusion gradients that develop as the biofilm thickens. Chavali and colleagues recently showed that a heterotrophic initial growth phase leading to electrode coverage would be necessary for cathodic growth of *C. necator* to proceed (Chavali *et al*., 2026). The rationale is that dense electrode coverage leads to biotic respiratory oxygen depletion before it can reach the cathode surface, thereby suppressing ROS formation. Therefore, under oxic conditions, close contact ceases to be merely advantageous and becomes a prerequisite. Still, this phase also took a considerable amount of time and general solutions to enhance or accelerate growth on electrodes are so far missing. No systematic engineering or selection strategy has been reported to specifically improve its biofilm formation and performance on cathodes, even though its evolvability and its electroautotrophic potential have each been demonstrated in isolation.

Adaptive laboratory evolution (ALE) offers a direct route to close this gap. By cultivating populations under defined selective pressures over many generations, ALE allows spontaneous, fitness-enhancing mutations to accumulate and become fixed without prior knowledge of their targets (Mavrommati *et al*., 2022). In *C. necator*, ALE has already improved tolerance to inhibitory gases such as carbon monoxide under both heterotrophic and autotrophic conditions, with genome sequencing of evolved isolates pinpointing causal mutations in respiratory and hydrogenase genes (Wickham-Smith *et al*., 2023). ALE was also used for *C. necator* in bioelectrochemical systems based on planktonic *C. necator* cultures in which the organism was adapted to higher salt concentrations which reduced ohmic losses in the system (van der Sande *et al*., 2026). In parallel, selection-based approaches in other MES model organisms have yielded cathodic strains with markedly improved biofilm accumulation, electrode coverage and resistance to electrochemically generated stress as mentioned above for *K. spormannii* (Jung *et al*., 2020), supporting the expectation that comparable gains are attainable for *C. necator*.

Here, we apply ALE to enhance electroautotrophic biofilm formation of *C. necator* on cathodes, aiming for strains that combine robust autotrophic growth with improved electrode colonization under oMES conditions. Populations were subjected to long-term cathodic, CO₂-based selection in a bioelectrochemical reactor; evolved isolates were genome-sequenced to link mutations to improved biofilm performance, and the most dominant isolate was phenotypically compared to the progenitor. A defined consortium of evolved strains was then screened in a microfluidic oMES platform to identify the strain that was subsequently validated in pure culture with quantitative monitoring of biofilm development. Together, these experiments test whether ALE can serve as a practical strategy to engineer *C. necator* for robust electroautotrophic biofilm performance and thus advance toward scalable oMES-based CO₂ valorization.

## 2. Materials and Methods

### 2.1 Adaptive laboratory evolution of *C. necator* electroautotrophic biofilm variants

The progenitor strain for this adaptive laboratory evolution was *Cupriavidus necator* H16 strain JG2015 (Δ*acoABC* Δ*phaC1/C2* Δ*adh*), which carries the empty plasmid pBBR1MCS-2. Using this strain enabled us to add 250 µg mL^-1^ of kanamycin throughout the process and thereby reduce the risk of contamination. A preculture of progenitor cells was grown in LB media (30 mL) at 30 °C with 180 rpm shaking for 36 to 48 hrs. The cells were then washed three times with 6000 rcf centrifugation in minimal media (Sydow *et al*., 2017) without an added carbon source and introduced into the flow-cell reactor (Hackbarth *et al*., 2020) filled with the same fresh minimal media (pH 6.83) to a final OD of 0.3-0.4. The three-electrode setup flow-cell reactor was operated at atmospheric pressure, with Viton tubing connecting to a reservoir bottle (Schott 500 mL with customized lid stopper) kept in a 32 °C water bath. The cathode, a 20 cm^2^ graphite electrode was polarised at −0.5 mA cm^-2^ by connecting the setup to a potentiostat (Interface 1010E, Gamry Instruments, Warmister, USA) with a reference electrode (SE23I, Sensortechnik Meinsberg, Waldheim, Germany) installed at the inlet of the flow cell. The six counter electrodes were positioned opposite the working electrode, with their cumulative contact surface equal to that of the cathode and they were made up of iridium-tantalum-coated titanium (Platinode™ – MMO Anode 177, Umicore, Schwäbisch Gmünd, Germany). A liquid flow rate of 20 mL min^-1^ (Ismatec ISM901B Compact digital console drive with remote control, 115 to 230 VAC, 60/50 Hz) was maintained throughout the operation of the flow-cell reactor. The flow-cell reactor and the peripherals were gas-tight and approximated to 60 mL in liquid volume while the reservoir bottle contained 200 mL. Every 2 days, a gas mix composed of 1.2 Ln hr ^-1^ CO_2_ and 0.4 Ln hr ^-1^ O_2_ (Bronkhorst EL-Flow Select) was purged into the liquid phase of the reservoir bottle for 30 minutes while the headspace was connected to the atmosphere. The media was changed after 3 to 5 days of initial inoculation to minimize remaining planktonic cells. The flow-cell operation was stopped after biofilm growth stabilized on the cathode as indicated by OCT imaging (Ganymede II, Thorlabs, Daschau, Germany). At the end of each run, the bacterial communities were removed from the cathode surface by repeatedly pipetting directly onto the electrode. The recovered cells were subsequently serially diluted and plated on both LB and minimal medium agar, the latter supplied with (need to look up gas composition numbers) H2:CO2:O2 = 7:2:1 and CO_2_ as the sole carbon source, to assess growth. The resulting colonies were used to confirm the bacteria as *C. necator* via 16S rRNA sequencing (Mix2Seq, Eurofins), as well as to prepare cryogenic stocks in 25% glycerol to store at –80 °C. A total of 5 passages were conducted, with flow-cell reactors operated for 31, 6, 18, 23, and 20 days, respectively. gDNA from 29 isolates from the last generation were extracted (DNeasy PowerBiofilm, Qiagen, Carlsbad, USA) and analyzed (MinION, Oxford Nanopore, Oxford, UK) for genetic variations from JG2015.

### 2.2 Microfluidic Cultivation Strategy

The 29 isolates were cultivated in a microfluidic cultivation platform setup following the approach of Klein and colleagues (Klein *et al*., 2024). The system was tailored to function as a cathodic microbial electrosynthesis system by maintaining it at a constant current of – 45.45 µA cm^-2^ using a potentiostat (PalmSens MultiEmStat4 LR). A dual-phase growth approach with an initial heterotrophic phase followed by electroautotrophic growth was chosen to allow for better attachment of cells onto the electrode surface. The growth media (Sydow *et al*., 2017), pumped at a steady volumetric flowrate of 4 mL hr^-1^, was supplemented with Fructose to act as a carbon source until the biovolume reached confluent electrode coverage, following which the carbon source was switched to gaseous carbon dioxide. After a total growth period of 14 days, the biomass was harvested for sequencing to identify the most abundant strain.

The same procedure was repeated with this isolate to study its performance and compare it to the progenitor and understand the influence of the adaption process.

### 2.3 Optical Coherence Tomography (OCT)

The mesoscopic structure of the biofilms during the adaptive evolution as well as microfluidic growth was examined by optical coherence tomography (OCT) as described by Klein et al., 2024. Images were acquired with a Ganymede™ spectral domain OCT system (GAN611C1-SP1, Thorlabs GmbH, Dachau, Germany), using the acquisition parameters reported previously (Bauer *et al*., 2019; Klein *et al*., 2024). The resulting datasets were analyzed in Fiji (Fiji Is Just ImageJ), version 2.1.0/1.53 (Schindelin *et al*., 2012). Biofilm characterization and quantification followed a sequence of processing steps. Biovolume was obtained by multiplying the number of signal-positive pixels by the voxel size of the image. Height maps of the cathodic surfaces were produced following the processing routine of Wagner and Horn (Wagner and Horn, 2017) and surface coverage was then derived from these maps.

Biofilm porosity was estimated by comparing the measured biofilm height (h_real_) with a theoretical height (h_theoretical_), calculated from the biovolume on the assumption of a fully dense (100%) biofilm, according to Equation 1 as follows:

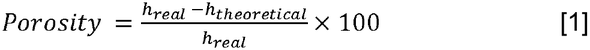

### 2.4 Sequencing

The 29 isolates were grown in 29 individual LB flasks and cells were harvested by centrifugation. The biofilm from the microfluidic chip experiment using a mixture of the 29 isolates was directly harvested from the electrode and centrifuged to obtain a cell pellet. DNA was isolated from the cell pellet using the Qiagen DNeasy Powerbiofilm kit according to the manufacturer’s instructions. DNA quality and yield were measured using a nanodrop 2000 and Qubit dsDNA HS Assay kit according to manufacturer’s instructions. Subsequently, DNA was prepared for sequencing using Oxford Nanopore Rapid barcoding kit 24 V14 based on the manufacturer’s instructions and sequenced on an PromethION P2 solo. Super-accurate base calling was performed with MinKNOW 25.05.15 with adapter and barcode trimming enabled.

Subsequent data analysis was performed using the CLC genomics workbench 20.0.4 (Qiagen). Sequencing reads from the progenitor strain JG2015, the 29 evolved isolates and the harvested biofilm community were mapped to the reference genome of *Cupriavidus necator* (ASM479872v1) and the pBBR1 plasmid sequence. Variants were called for all samples relative to ASM479872v1 using Fixed ploidy variant detection Initial thresholds for variant detection were modified to 30% required variant probability, minimum coverage set to 7, minimum count to 2 and minimum frequency to 10%. Comparative analysis of mutations and similarity between strains were done in Microsoft Excel. Variants already present in the re-sequenced progenitor strain were excluded (background subtraction) to distinguish evolution-derived variants from pre-existing strain-background differences. A pairwise comparison of mutations rates with minimum 50% frequency between each sample was used to create a distance table, which was subsequently used to identify similiarity based on a jaccard coefficent.

The annotated reference genome was screened for all genes with annotations containing “diguanylate cyclase”, “phosphoediesterase”, “GGDEF” or “EAL”. Further, translations of all gene sequences were searched for GGDEF motifs and variants with one or two substitutions. The overlap of both lists is summarized in Table 1.

**Table 1:**
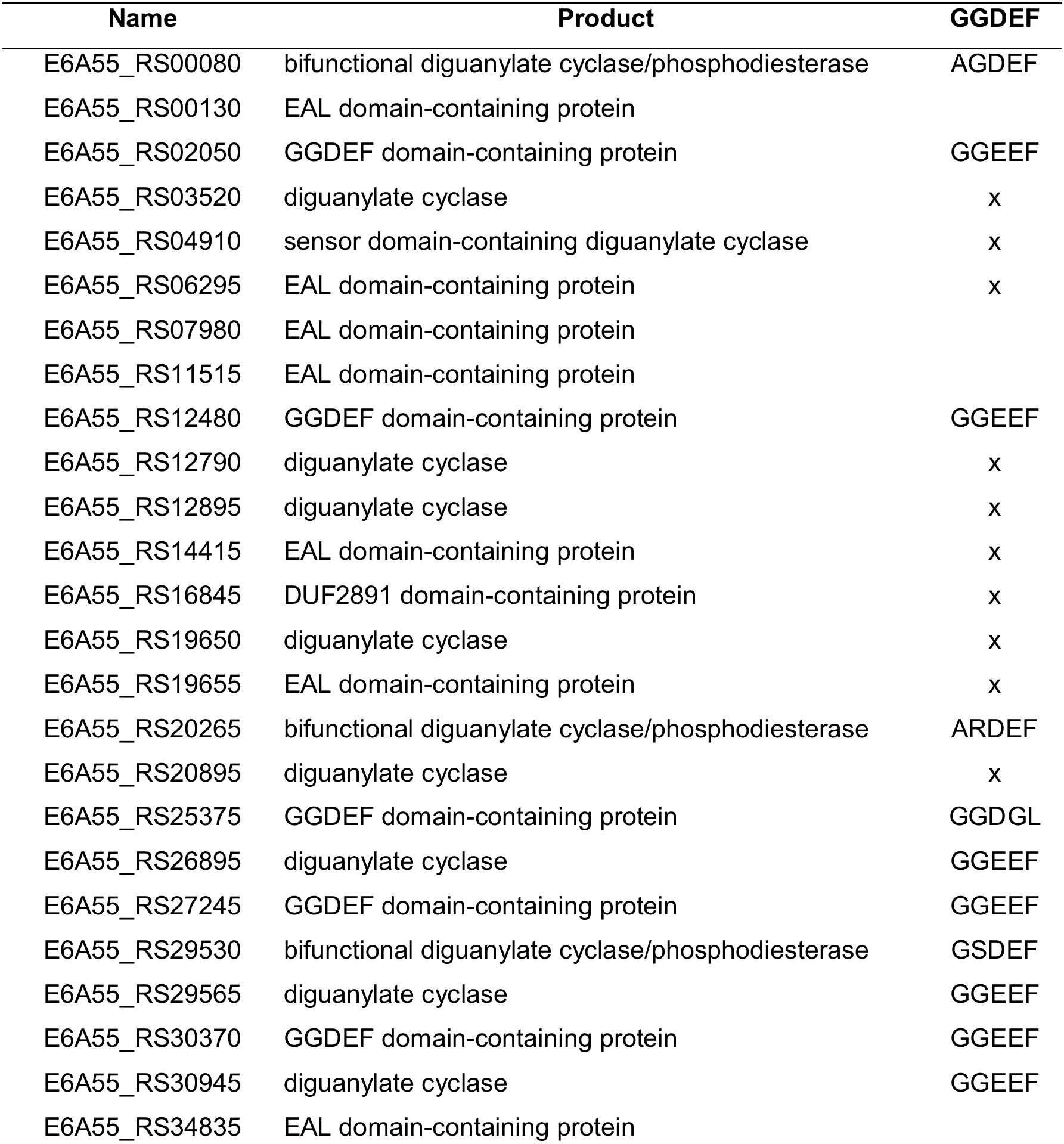
Candidate c-di-GMP signalling enzymes identified via genome annotation. Genes marked with an “x” carry an unaltered GGDEF motif.

The structural representation of the bifunctional diguanylate cyclase/phosphodiesterase and the resulting fragments of it, were prepared with PyMOL v3.1.8 (The PyMOL Molecular Graphics System, Version 3.0 Schrödinger, LLC.), with a model derived from AlphaFold server (Abramson *et al*., 2024). Coordination of substrates GTP and c-di-GMP were achieved by aligning structural orthologs with highest blastp (National Library of Medicine (US), 1988) similarity to the corresponding domains (3hv8, for EAL and 5m3c for GGDEF) in the PDB (Berman, 2000).

### 2.5 μ-Particle Tracking Velocimetry (μ-PTV)

The μ-PTV technique outlined in (Merbach *et al*., 2026) was applied to obtain time-resolved 3D particle trajectories within the microfluidic reactors.The setup comprised an optical system (μ-PTV system) and the microfluidic cultivation platform without biofilm. The optical system is part of the Major Research Instrumentation *MUST* (LaVision GmbH, Göttingen, Germany). The measurement system consists of a Nd:YLF dual-cavity laser providing a maximal pulse energy of 30 mJ per cavity at a maximum repetition rate of 10 kHz and an emission wavelength of 527 nm (Photonics Industries International Inc., USA), which was directed through a stereo microscope (SteREO Discovery.V12, Carl Zeiss AG, Germany), to illuminate the volume of interest in the microreactor. The resulting two stereo-views have been recorded by two high-speed cameras (Phantom VEO 640L, Vision Research, USA), the timing of illumination and recording is controlled by a synchronising timing unit. The laser output intensity was adjusted with the operating current and an integrated attenuator that were set to 30 A and 75% for both cavities. The laser excited fluorescent polymethyl methacrylate particles (PMMA-RhB-Fi89, ρ = 1.19 g cm^−3^, microParticles GmbH, Germany) with a diameter of *d*_p_ = 10.14 ± 0.26 μm were pumped through the microfluidic channel at 4 mL hr^−1^. The particles have a maximum excitation wavelength of 560 nm, and the band overlaps with the emission of the Nd:YLF laser, whereas their fluorescence maximum is located at 584 nm. The image acquisition frequency of the cameras was set to 25 Hz. Both high-speed cameras have a resolution of 2560 × 1600 pixels. They were arranged such that their optical axes were aligned with the direction of the incident laser beam. The microscope operates at magnifications in the range from 1.28-fold to 16-fold. The maximal numerical aperture is 0.03. The present experiments were performed at a total magnification of 2.5 such that the resulting Field Of View (FOV, width) is approx. (18 x 4) mm^2^ and the Depth Of Field (DOF), which is inversely proportional to the microscope magnification, is approx. 1.5 mm. The μ-PTV measurements are evaluated using DaVis 11.2 software (LaVision GmbH, Germany) for particle reconstruction and tracking using the variable time-step Shake-The-Box algorithm (Schanz *et al*., 2016). Furthermore, the actual channel profile, as obtained by profilometry (Figure 2), was used as a geometric mask during the particle tracking to suppress the detection of ghost particles. The detailed software and the image pre-processing settings are shared via the TORE repository.

### 2.6 Profilometry

To characterise the geometry of the microfluidic channel, the 3D Optical Profilometer VR-6000 (Keyence Deutschland GmbH, Germany) was used. Structured light was diagonally projected on the sample from two sides. Height variations on the surface cause a displacement of the projected light pattern in proportion to the surface topography. The distorted light pattern was captured by a camera from above the sample. To replicate the three-dimensional sample surface, the triangulation principle, more specifically, the optical sectioning method was applied to calculate the surface height from the known illumination angle relative to the optical axis of the camera and the measured distortion of the structured light.

For the measurement, the microfluidic cultivation platform was placed on the sample holder in a way that allowed maximum illumination of the channel from the two LED light sources. Images and surface data were recorded at a magnification of 40× and automatically stitched together to capture the whole geometry. The obtained surface profile was exported as STL file, which was then used in the evaluation of the µ-PTV measurements.

## 3. Results and Discussion

### 3.1 Progenitor growth behaviour

The aim of this study was to use ALE to advance the growth of *C. necator* on cathodes*(Reiner et al., 2020)*because under purely autotrophic conditions, *C. necator* did not colonise the graphite electrodes directly. Instead, robust growth was only obtained when a confluent coverage of the electrode was first established during an initial heterotrophic phase (Chavali *et al*., 2026). This requirement is most likely a consequence of abiotic oxygen reduction at the cathode, which generates reactive oxygen species (ROS) that impair initial cell attachment and early biofilm development. To this end Chen et al. reported previously that growth of *C. necator* in cathode compartments can be increased by developing strains displaying superoxide dimutase on the cell surface ((Chen et al., 2023). This study was conducted with planktonic cells and even here, although the local concentration of ROS might likely be lower compared to cathode-biofilm systems, growth could be accelerated more than twofold (Chen et al., 2023). Under heterotrophic conditions, six days were required to reach confluent electrode coverage (Figure 1). This is a rather long time considering that in these six days the biofilm was not productive regarding CO_2_-based biomass production. Still, this requirement of heterotrophic on growth was also described for anaerobic systems, where it was revealed that heterotrophic on-growth could be more beneficial for acetogenic productivity compared to an initial phase of voltage removal (Zhang *et al*., 2022).

**Figure 1:**
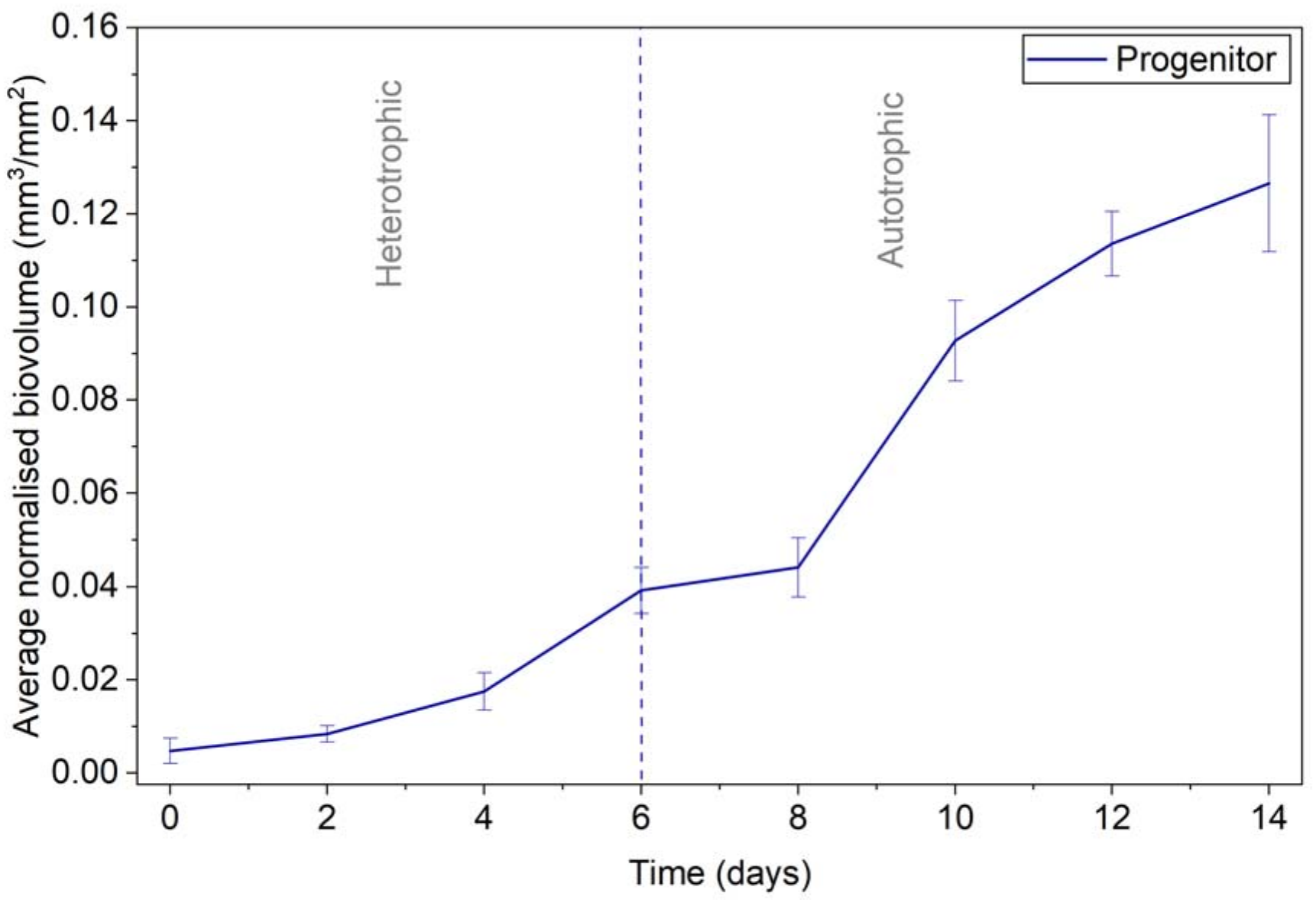
Biofilm development by *Cupriavidus necator* progenitor strain JG2015 on a graphite electrode within a microfluidic oxic microbial electrosynthesis (oMES) system under biphasic growth conditions. Data points represent the average of triplicate measurements (n = 3 independent reactors operated in parallel).

**Figure 2:**
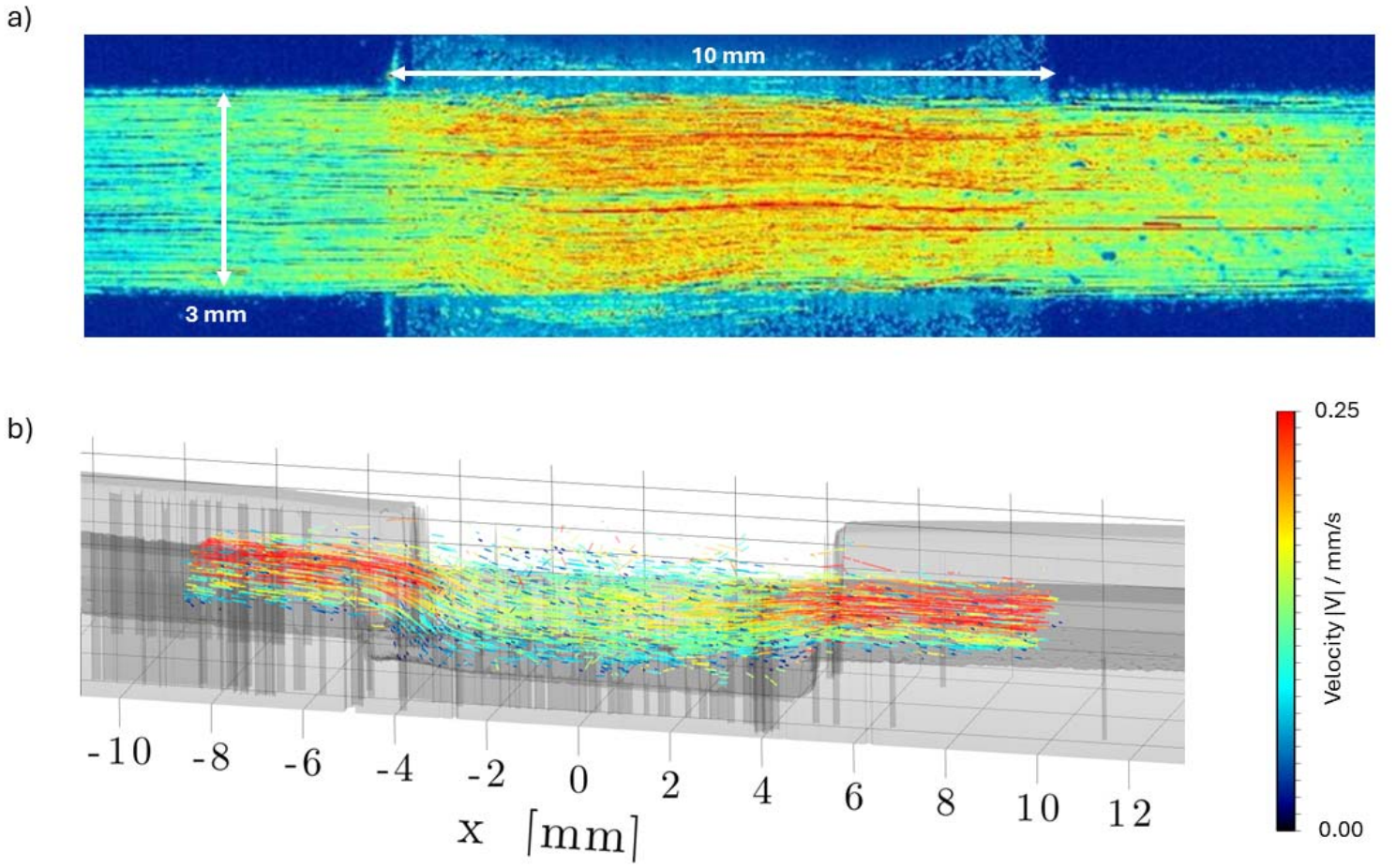
Flow through the microfluidic cultivation platform without biofilm from left to right. a) Time-averaged image of the flowing fluorescent particles of 10 µm size colored according to the maximum– minimum counts at each pixel, such that yellow to red areas show that more particles have been passing through the corresponding pixel. b) 3D and time resolved particle trajectories in the channel profile, coloured according to their absolute velocity |V| and displayed with a track length of 200 images, representing 8 s, f = 25 Hz.

After the switch to autotrophic growth (with CO₂ as the sole carbon source), the biofilm biovolume increased from 0.04 to 0.12 mm³ mm⁻² within eight days. However, this growth was not uniform across the flow channel: thicker biofilm developed only in the downstream part of the electrode, whereas the upstream part showed patchy colonisation, with almost no growth at the leading upstream edge. To identify the cause of this spatially heterogeneous colonisation, we characterised the local flow conditions above the electrode using micro-particle tracking velocimetry (μ-PTV).

### 3.2 Biofilm morphology and influence of flow characteristics

µ-PTV revealed a pronounced reorganization of the flow field across the electrode (x = −5 to +5 mm), driven by the local increase in channel height in this region (Figure 2). In the narrow inlet channel the flow reached up to ∼0.25 mm s⁻¹, but at the upstream (leading) edge of the electrode (x = −5 mm) the sudden expansion of the channel cross-section caused this fast near-wall flow to separate: the high-velocity streamlines plunged from the upper channel toward the electrode surface, while a zone of low, disorganized velocity, a recirculation vortex developed behind the expansion step and above the upstream half of the electrode (Figure 2 b). The reconstructed channel geometry obtained by profilometry confirmed the topographical step at this junction as the origin of the swirl. Over the downstream half of the electrode (x ≈ +1 to +5 mm) the flow reattached and re-developed into a more continuous, uniform stream, with full recovery of the bulk channel profile only beyond the trailing edge (x > +5 mm).

This flow structure can provide a coherent explanation for the heterogeneous colonisation described above. In the upstream recirculation zone, the unsteady flow could hinder stable cell attachment and early biofilm establishment, consistent with the patchy growth and the near-absence of biofilm at the leading edge. Literature is rich in showing the impact of flow on biofilm architecture. A coupling between flow separation at geometric edges and localised biofilm deposition has been demonstrated, for instance, for flow around corners and through porous materials, where secondary flows trigger directed biofilm and streamer formation (Rusconi *et al*., 2010; Drescher *et al*., 2013; Tsagkari *et al*., 2022). Over the downstream half, where the flow reattaches and becomes uniform and low in shear, cells attach and develop into a confluent biofilm.

Because the flow field was measured in the absence of biofilm, its influence on biofilm architecture is necessarily inferred; nevertheless, the endpoint morphology mirrors the flow structure. OCT B-scans and height maps of the mature biofilm (Figure 3) show taller, pillar-like structures interspersed with shorter sections in the entrance region, which give way to a highly uniform biofilm toward the outlet. This spatial pattern is consistent with the established observation that variable or elevated shear drives a morphological transition toward raised bio-aggregates, streamers and pillars, whereas stable, low-shear environments favour even, uninterrupted growth. (Tsagkari and Sloan, 2018; Lee *et al*., 2023)

**Figure 3:**
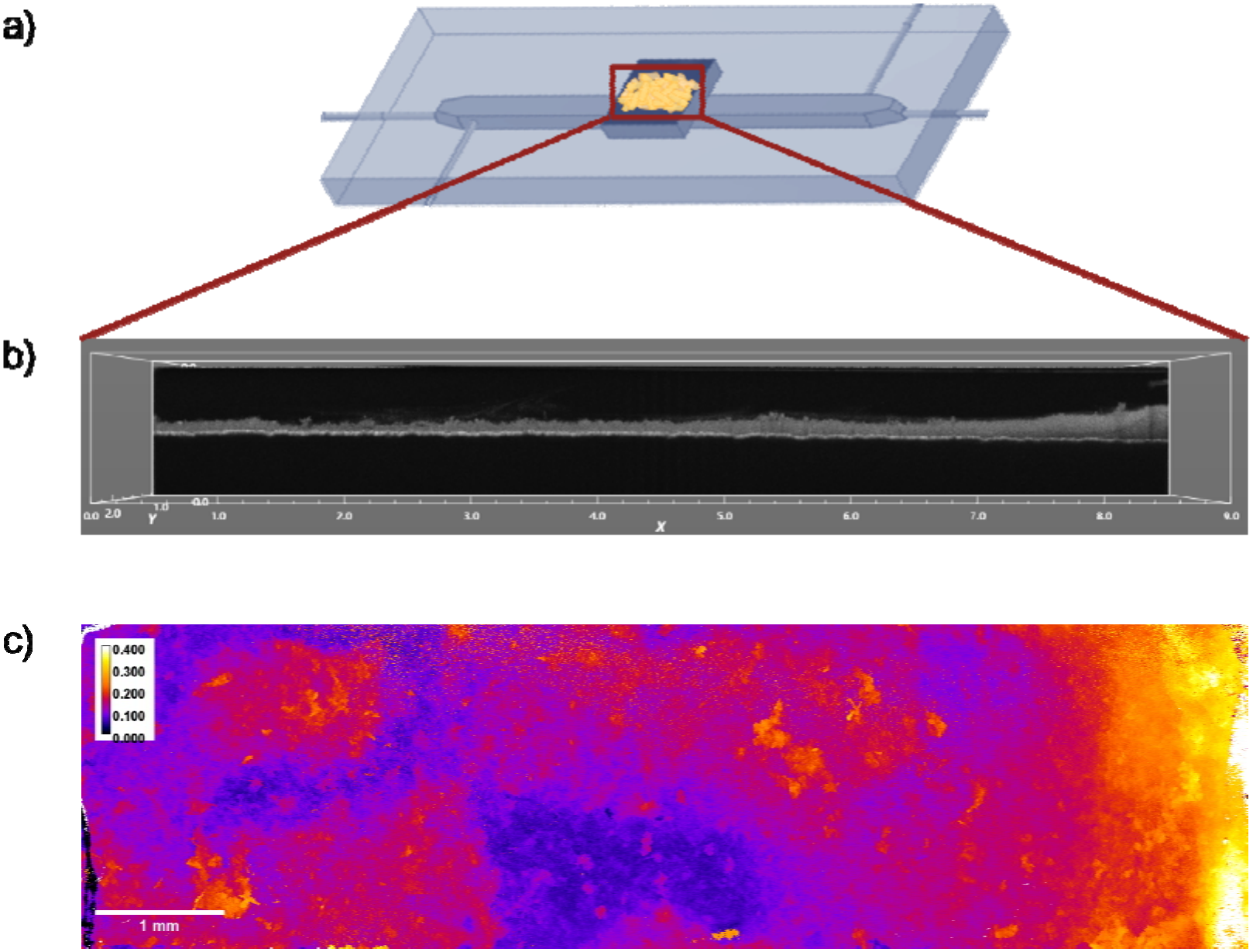
Three-dimensional structural visualization and height variation of the *Cupriavidus necator* progenitor biofilm along the length of the electrode after 14 days of growth. (a) Schematic representation of a microfluidic reactor with electroactive biofilm growth on graphite electrode (b) Optical coherence tomography (OCT) B-scan displaying a representative x-z cross-section of the biofilm on the electrode. (c) Endpoint height map illustrating the spatial variance in biofilm thickness in millimeters.

Beyond this mechanical effect, the recirculation zone may act through a second, mass-transfer-related mechanism. As this zone is poorly exchanged with the bulk liquid, local convective transport of O₂ is reduced and reaction products accumulate; such mass-transfer limitation is known to govern both the performance and the spatial distribution of aerobic biocathode biofilms (Milner and Yu, 2018). At the same time, the observed recirculation might lead to an accumulation of ROS, which would lead to an increased localised inhibition of biofilm growth.

### 3.3 Adaptive laboratory evolution and population structure

For the evolution experiments we used a larger bioelectrochemical flow cell (Reiner et al., 2020) initially, which allowed each passage to be performed with substantially higher cell numbers than the microfluidic platform. In this setup the oxygen input was actively controlled, which is expected to limit abiotic ROS generation at the cathode; under these conditions the initial heterotrophic on-growth phase could be omitted and biofilms established directly under autotrophic conditions. Once biofilm growth had levelled off into a steady state, the reactor was opened and a dilution series was plated onto agar plates. Clones from agar plates were pooled and used for reinoculation. Adaptation was carried out over five successive passages.

Following five cycles of adaptation, the adapted strains demonstrated a distinct shift in growth behavior and a significantly enhanced capacity for biofilm formation compared to the progenitor strain. Over the 15-day experimental period, the progenitor strain maintained a consistently low biovolume, with a final yield of approximately 0.01 mm^3^ mm^-2^. In contrast, the post-adaptation strains exhibited rapid initial accumulation, reaching an early peak biovolume of over 0.08 mm^3^ mm^-2^ by day 4. Although this initial surge was followed by a reduction, the adapted strains maintained a consistently higher baseline throughout the remainder of the experiment, ultimately yielding a final biovolume of roughly 0.04 - 0.045 mm^3^ mm^-2^ on day 15.

Because clones were pooled from the agar plates rather than isolated individually, it was initially unclear whether selection had converged on a single dominant genotype or whether the biofilm comprised several clones with distinct mutational profiles. To resolve this, we picked 29 random colonies after the 5th passage, expanded them in liquid culture and sequenced their genomes. Variants were called against the reference genome and compared to the sequenced progenitor JG2015, so that fixed strain-background differences were excluded and only mutations acquired during the evolution experiment were counted. When applying stringent filtering thresholds (min. 90% allele frequency), the resulting mutation profiles of the liquid-grown isolates could not account for the high-frequency mutations later found in the mixed biofilm, as even key adaptive mutations (such as E6A55_RS29530) occurred at frequencies below 90%, possibly due to partial reversion or counter-selection during planktonic growth. We therefore performed an initial permissive variant calling in CLC Genomics Workbench followed by cross-sample filtering in Excel to capture sub-clonal variants shared with the mixed biofilm. Under these relaxed criteria, each liquid-expanded isolate exhibited a broad profile of 156–352 predominantly low- and intermediate-frequency variants, which we used to assess similarity to the cathode biofilm community. The population recovered from the cathode was therefore not a single evolved genotype but a mixture of co-existing lineages, each defined by its own mutational profile. Such coexistence of multiple adapted lineages within one evolving population is a hallmark of clonal interference and has been observed across microbial evolution experiments (Lang et al., 2013).

To assess the collective phenotype of this evolved population, the 29 isolates were grown individually, mixed in equal cell numbers, and the pool was used to inoculate the microfluidic flow cell for direct comparison with the progenitor. Growth behavior changed markedly compared to the progenitor strain shown in Fig.1. Confluent biofilm coverage was reached after only one day of heterotrophic growth, compared with six days for the progenitor. Under subsequent autotrophic conditions, the biovolume first rose to 0.14 mm³ mm⁻² and then declined to a steady-state value of 0.06 mm³ mm⁻². The evolved pool thus accelerated initial electrode colonization and reached a higher peak biovolume than the progenitor (∼0.12 mm³ mm⁻²), but settled at a lower steady-state biovolume.

We interpret this behavior primarily as a trade-off between colonization speed and biofilm robustness. Selection in our system acted on rapid attachment and spreading, but the extracellular polymeric matrix that provides a biofilm’s cohesion and its adhesion to the substratum is metabolically costly to produce (Xavier and Foster, 2007; Flemming and Wingender, 2010). Strains selected for fast colonization may not at the same time produce a mechanically robust biofilm matrix. In this case, the weaker biofilm might be more readily removed under the local hydrodynamic shear characterized in Section 3.2. This characteristic of different EPS productivities and maybe chemistries might also be a property of individually adapted strains (fast colonizers versus slow robust biofilm builders). A further mechanism that may act in parallel is an active escape response that lowers standing biomass once the biofilm is established and as soon as resources become limited due to mass transfer limitation (Kaplan, 2010). Distinguishing these contributions would require direct measurement of matrix composition and of planktonic cell release during the decline phase, which was beyond the scope of this study. However, it is important to consider the complex cultivation history of the adapted strains, which underwent sequential shifts between heterotrophic planktonic pre-cultures, electroautotrophic biofilms, agar-based heterotrophic or autotrophic growth, and subsequent heterotrophic planktonic phases. By specifically harvesting cells that remained adhered to the cathode post-disassembly, this methodological approach may have unintentionally biased selection toward rapid colonizers. Such phenotypes demonstrate high plasticity in shifting from heterotrophic suspension to electroautotrophic biofilms, a metabolic transition selected for irrespective of the total final biofilm yield.

A noteworthy aspect is the apparent change in porosity over the course of ALE. Initially, the progenitor strain formed a comparatively dense biofilm with a porosity of 43.23%. However, after the series of evolutionary cycles designed to improve performance, the mixture of the 29 isolates produced a biofilm that at day 14 showed a porosity of 75.70%. This looser, open structure might be advantageous to overcome mass transfer limitations.

To determine which of the 29 isolates prevailed under cathodic conditions, the cathode biofilm was harvested at the end of the experiment and the community was re-sequenced. Because extensive variants with varying frequencies overlap among the isolates precluded a simple deconvolution via strictly private marker mutations, we calculated the genetic similarity between each isolate and the mature biofilm community using the Jaccard similarity index (and pairwise variant-frequency comparisons) across all called variant positions (Figure 5). Strain 890 showed the highest similarity to the harvested biofilm community, differing in only 29 variant positions (present in the biofilm at an average variant allele frequency of 60%, maximum 89%), followed by strain 921, which differed in 39 positions (average variant allele frequency of 77%, maximum 94%).

**Figure 4:**
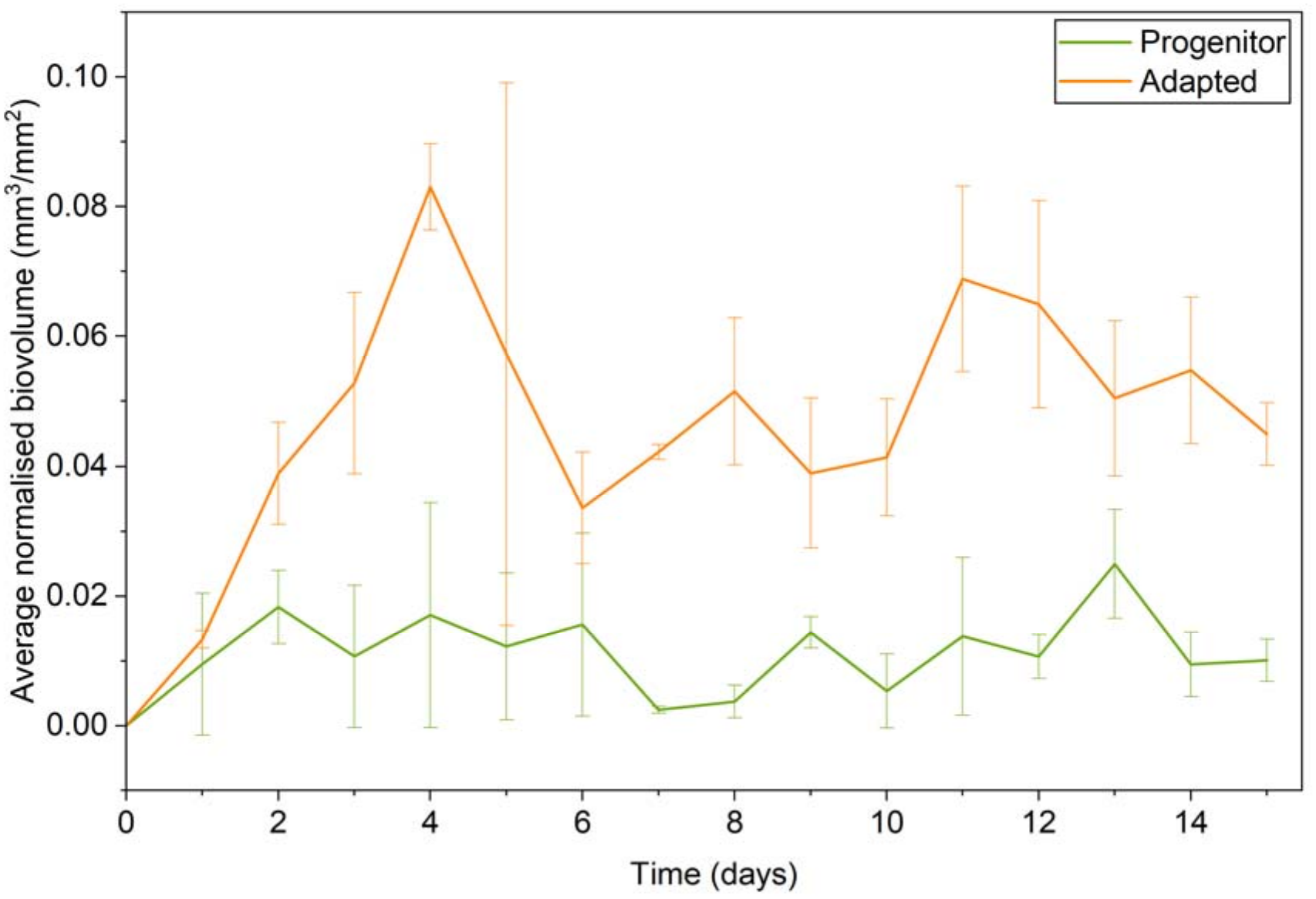
Biofilm development by *Cupriavidus necator* progenitor strain JG2015 in comparison to the adapted strains after 5 passages in a flow cell system under electroautotrophic growth conditions. Data points represent the average of duplicate measurements (n = 2 independent reactors operated in parallel).

**Figure 5:**
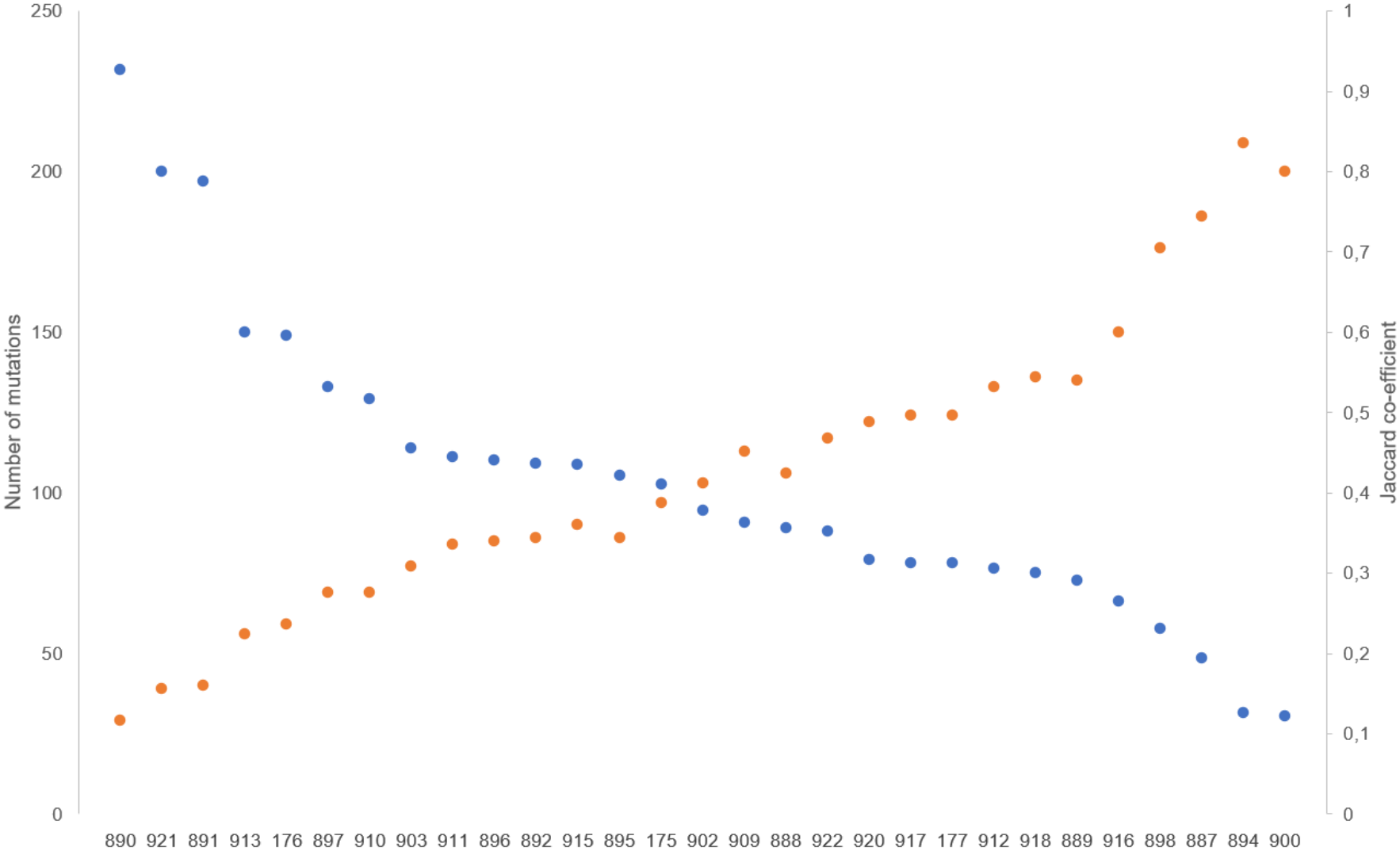
Genetic similarity between the 29 evolved *Cupriavidus necator* isolates and the harvested mixed-culture biofilm community from the microfluidic cultivation platform, quantified via differing variants and derived Jaccard similarity index.

### 3.4 Quantitative comparison of *Cupriavidus necator* biofilm in microfluidic cultivation platform: Progenitor vs Adapted strain

To test whether the pure strain 890 would behave differently from the mixed community of 29 isolates, we repeated the microfluidic bioelectrochemical experiment using strain 890 alone as the inoculum. As with the strain mixture, confluent coverage was again reached within only one day of heterotrophic growth, confirming that accelerated initial on-growth is a robust, heritable trait of the adapted strain rather than a property of the multi-strain community. The subsequent behaviour, however, differed from that of the mixture: under autotrophic conditions the biovolume increased steadily until it reached approximately the same value as the progenitor (∼0.12 mm³ mm⁻²) and then remained stable over the last four days of the experiment (Figure 6). In contrast to the mixed pool, which overshot to a higher peak biovolume (0.14 mm³ mm_⁻_²) before declining to a lower steady state (0.06 mm³ mm_⁻_²), strain 890 reached a lower peak but formed a more robust, stable mature biofilm. Thus, the multi-strain inoculum appears to promote a higher transient peak at the expense of long-term biofilm stability, whereas the accelerated heterotrophic on-growth is common to both and independent of community composition. Coming back to the hypothesis of differently adapted cell lines mentioned in section 3.3, strain 890 might be a more robust but slower biofilm builder compared to other evolved strains that in coculture dominate initial phases of biofilm development. To this end future experiments will comprise time resolved analysis of the percentage of individual strains over time. Strain 890 produced a biofilm characterized by a porosity of 79%, which is the highest porosity measured in all biofilm experiments conducted in this study. Again, this porosity might aid in overcome mass transfer limitation.

**Figure 6:**
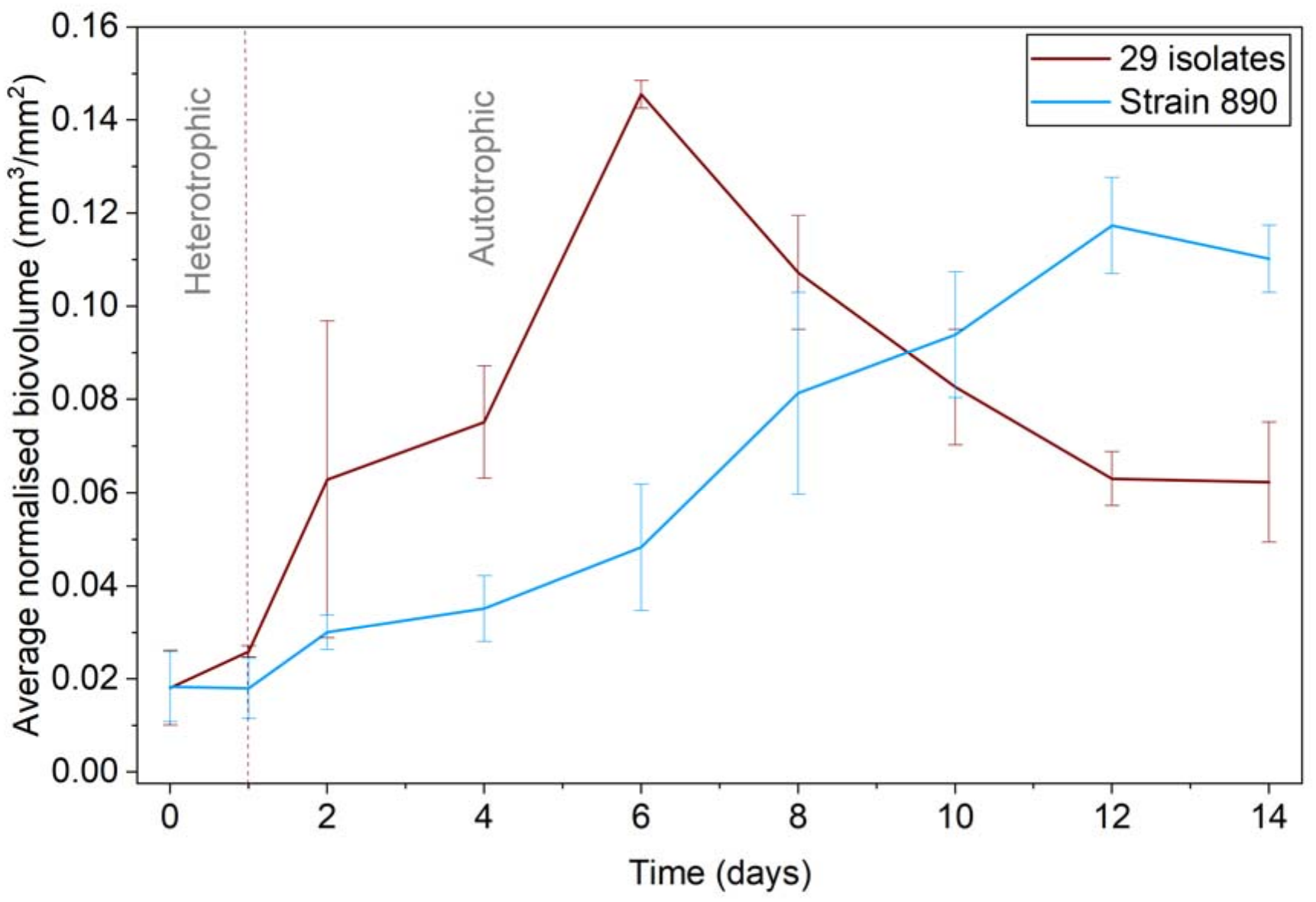
Comparison of biofilm development of an evolved 29-isolates *Cupriavidus necator* consortium to that of the dominant isolate (Strain 890) on graphite electrodes within a microfluidic oxic microbial electrosynthesis (oMES) system under biphasic growth conditions. Data represent the average of triplicate measurements (n=3 independent reactors operated parallely). The transition from heterotrophic to autotrophic growth was initiated after 1 day (upon reaching an initial surface coverage range of 0.01-0.03 mm^3^ mm^-2^), in contrast to 6 days needed for the progenitor strain.

The observation that the steady-state biovolume of the adapted strain did not exceed that of the progenitor is expected from the supply constraints of electroautotrophic Knallgas growth. The electron donor (H₂, generated at the cathode surface) and the terminal electron acceptor (O₂, supplied from the bulk liquid) enter the biofilm from opposite sides and establish counter-directional gradients, so that sustained growth is confined to the biofilm region in which both are simultaneously available (Rittmann, 2018). This co-limitation imposes a mass-transfer-limited ceiling on the steady-state biovolume that is largely decoupled from the traits selected during adaptation. Raising this ceiling would require simultaneously enhancing the uptake of both H₂ and O₂, a combination unlikely to arise from the selection applied here.

### 3.5 Genomic basis of accelerated biofilm formation: strain 890 versus the progenitor lineage

To pinpoint the genetic changes responsible for the accelerated cathodic biofilm formation of strain 890, we resequenced its mature biofilm and compared the variant profile with the variant profile of the mixed biofilm and a progenitor biofilm grown under identical conditions. Against a shared background of several hundred pre-existing variants, the two biofilms diverged surprisingly at only one non-silent, high frequent locus that was absent in the progenitor strain. This locus is the gene for an annotated bifunctional diguanylate cyclase/phosphodiesterase (E6A55_RS29530). These enzymes would typically carry a GGDEF (typically a diguanylate cyclase activity) and EAL (typically a phosphodiesterase domain) domains. These opposing activities synthesise and hydrolyse bis-(3′→5′)-cyclic dimeric GMP (c-di-GMP), a second messenger that serves in many microorganisms as the master switch of the bacterial biofilm lifestyle (Hengge, 2009; Schirmer and Jenal, 2009; Römling *et al*., 2013; Jenal *et al*., 2017). High intracellular c-di-GMP tips cells toward sessility, adhesion and matrix secretion while low levels favour motility and dispersal. Nevertheless, closer inspection of the progenitor sequence revealed that the corresponding protein carries a GSDEF instead of a GGDEF motif. For *Thermosynechococcus elongatus* it was shown that this sequence variation in a protein also carrying an EAL domain led to a GTP concentration dependent activity modulation of the phosphodiesterase domain (Enomoto *et al*., 2015). Hence, the GSDEF-domain acts as a GTP sensor rather than a cyclase. Strain 890 carries an 8 nucleotide tandem duplication in this gene. This duplication leads to a new stop codon after amino acid residue 225 so that the protein is now split in two open reading frames (Figure 7). Figure 7 (a) shows the corresponding gene sequence with translations. The first part of the gene contains the N-terminus that would most probably be still expressed by *C. necator*.

**Figure 7:**
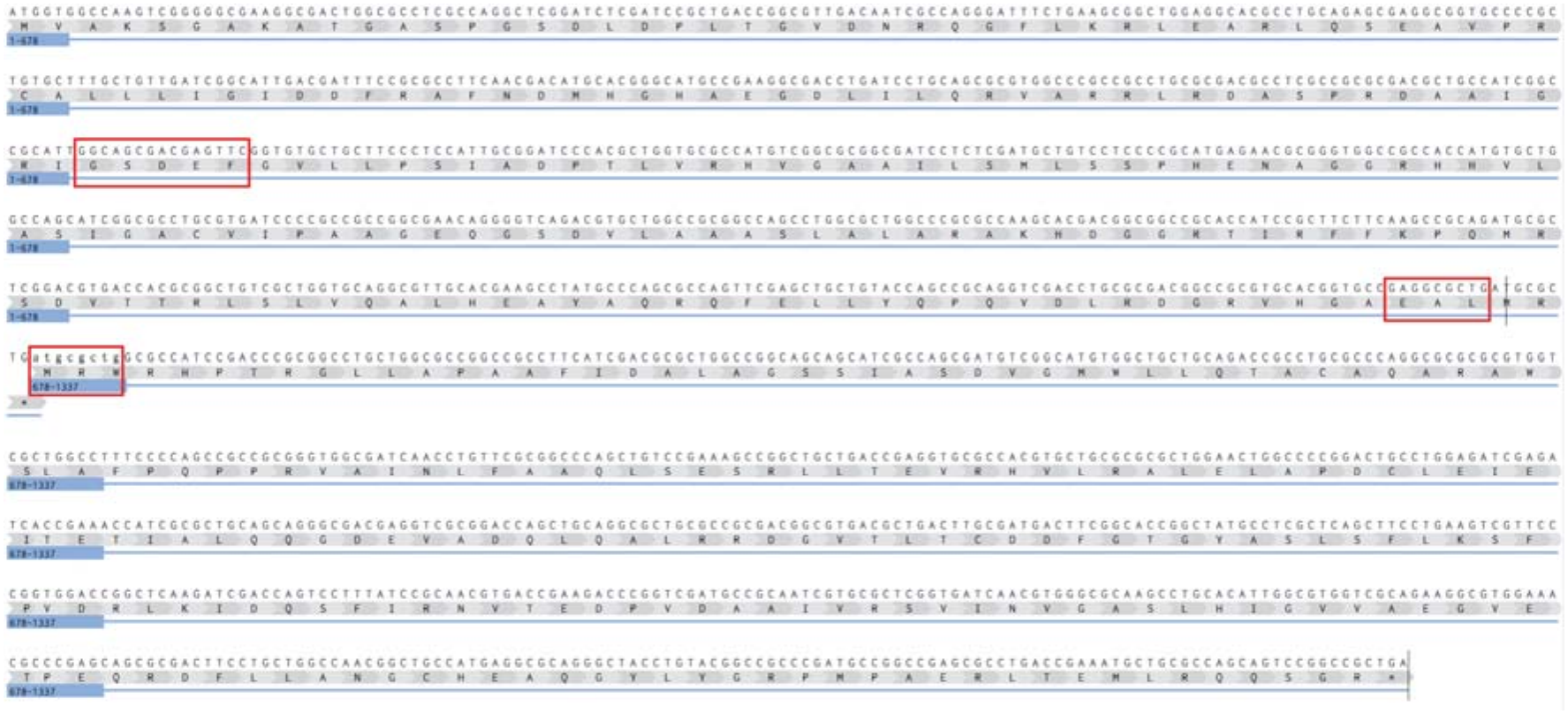
Sequence of bifunctional diguanylate cyclase/phosphodiesterase(E6A55_RS29530) with GSDEF, EAL motif and 8 basepairs duplication in position 678 hihglighted in red boxes Representation of sequence and annotation were done in benchling (Benchling [Biology Software]. (2026). Retrieved from https://benchling.com).

This part contains the GSDEF domain which would usually be connected to the EAL domain by an alpha-helix. The new stop codon is inserted two amino acids after the EAL domain. Hence, the EAL domain is now disconnected to the substrate binding pocket. Therefore, given the strongly biofilm-forward phenotype of strain 890, a plausible expectation is that the mutations caused a shift toward a net higher c-di-GMP concentration by eliminating the phosphodiesterase activity. Although the phenotype fits excellently to the observed mutation, future experiments will still be conducted in order to quantify c-di-GMP concentration in strain 890 compared to the progenitor.

Certainly E6A55_RS29530 is not the only enzyme in *C. necator* that could modulate the intracellular c-di-GMP concentration. The annotated genome lists 25 candidate enzymes with diguanylate cyclase or phosphodiesterase activity or being bifunctional. Closer inspection reveals that nine of these proteins contain a canonical GGDEF motif while we identified 7 proteins containing an GGEEF and with E6A55_RS29530 only one containing a GSDEF motif. While a GGEEF sequence was shown several times to be combined with a diguanylate cyclase activity (Christen *et al*., 2005), GSDEF was as mentioned before for one organism revealed as sequence variation leading to a GTP dependent activation of the phosphodiesterase activity. Interestingly, a similar case was discovered for the sequence variation GEDEF in *Caulobacter crescentus*. Also, here a variation in the second amino acid led to GTP sensor domain that would modulate the phosphodiesterase activity of the adjacent EAL domain (Christen *et al*., 2005).

Recently, Weiler et al. published results from a transposon screen aiming to discover mutations that would enhance biofilm growth under heterotrophic conditions (Weiler *et al*., 2026). The authors discovered several mutations leading to increased biofilm. Amongst these mutations, was also the disruption of E6A55_RS29530. When the authors tested a clean deletion of the gene they observed not faster biofilm growth kinetics but faster on-growth which would most likely be due to better initial colonization behaviour. Exactly the same could be shown here by the faster heterotrophic colonization of the cathode surface. Hence, the apparent most competitive solution for cathodic growth is not specific to electroautotrophic growth conditions. This is fundamentally different to a previous study conducted with *K. spormannii* in which ALE for electroautotrophic growth led to mutations that were likely combined with higher resilience against oxidative stress as well as with an impact on sulphur metabolism (Jung *et al*., 2020).

## 4. Conclusion

This study establishes adaptive laboratory evolution as an effective strategy to improve the electroautotrophic biofilm performance of *C. necator* on cathodes, and it links the resulting phenotype to a remarkably compact set of candidate mutations. The progenitor colonised graphite electrodes only after heterotrophic priming and formed spatially uneven biofilms whose distribution might be impacted by the local hydrodynamics. Evolution under controlled, oxygen-limited cathodic conditions relaxed these constraints and yielded a genetically heterogeneous population, from which the isolated strain 890 reached the target coverage in one rather than six days and maintained a robust biofilm at the progenitor’s steady-state biovolume.

The two principal barriers to cathodic colonisation were addressed by the same strategy. The kinetics of initial attachment were overcome genetically: strain 890 differed from the progenitor at only at only one locus, encoding a potential central regulators of the planktonic-to-biofilm transition. Oxidative stress, was likely mitigated by fast oxygen depletion catalysed by the developed strain. The steady-state biovolume remained close to the progenitor level, consistent with a mass-transfer ceiling imposed by the counter-directional supply of hydrogen from the cathode and oxygen from the bulk, which confines active growth to the region where both are simultaneously available and is largely decoupled from the traits under selection.

Together, these results show that the initial-attachment barrier to cathodic colonisation is accessible to evolution and associated with a small, tractable mutation, whereas steady-state productivity remains bounded by the physics of substrate co-supply. Raising the productivity ceiling will require enhancing hydrogen and oxygen uptake simultaneously, which is a combination unlikely to arise from the selection applied here. For oxic microbial electrosynthesis, where rapid, uniform coverage and a stable, harvestable biofilm govern space–time yield, strain 890 nonetheless represents a robust chassis and a well-defined starting point for CO₂ valorisation.

## Acknowledgements

The flow measurements were performed using the Major Research Instrumentation MUST (mobile multi-analysis unit for flow investigations in the context of power-to-x technologies and hydrogen applications), funded by the DFG (514139948) (equipment DOI: 10.15480/882.16713).

The geometry of the microfluidic channel was characterized using a 3D Optical Profilometer VR-6000 (Keyence Deutschland GmbH, Germany) (equipment DOI: 10.15480/882.14796).

## Data availability statement

All supporting data collected and reported in this study will be made available at TUHH Open Research (TORE) - https://doi.org/10.15480/882.17914

The raw sequencing data generated during this study have been deposited in the NCBI Sequence Read Archive (SRA) under accession number PRJNA1510941.

